# Decoding the nonconscious dynamics of thought generation

**DOI:** 10.1101/090712

**Authors:** Roger Koenig-Robert, Joel Pearson

## Abstract

Much of economics, psychology and neuroscience have focused on thought dynamics and how they control our behavior, from individual moral choices to the irrationality of market dynamics. However, how much of our thoughts we actually control when we feel we make deliberate choices remains unknown. Here we show that the content of thoughts can be decoded from activity patterns as early as 11 seconds before individuals report having formed the volitional thought. Participants freely chose which of two differently oriented and colored gratings to think about. Using functional magnetic resonance imaging (fMRI) and pattern classification methods, we consistently classified the contents of thoughts using activity patterns recorded before and after the thought was reported. We found that activity patterns were predictive as far as 11 seconds before the conscious thought, in visual, frontal and subcortical areas. These predictive patterns contained similar information to the responses evoked by unattended perceptual gratings and were evident in individual visual areas. Interestingly, neural information present before the decision was associated with the vividness of future thoughts, suggesting that preceding nonconscious sensory-like representations can impact the content and strength of future conscious thoughts. Our results suggest that thoughts and their strength can be biased by prior spontaneous nonconscious perception-like representations, advancing theories of free will and models of intrusive and repetitive thought production.

---

A large amount of psychology and, more recently, neuroscience has been dedicated to examining the origins, dynamics and categories of thoughts (*1*–*3*). Sometimes thoughts feel spontaneous and even surprising, while other times they feel effortful, controlled and goal orientated. When we decide to think about something, how much of that thought do we actually control?

To investigate this question, we crafted a thought-based mental imagery decision task, in which individuals could freely decide what to think about, while we recorded brain activation using fMRI. This paradigm enabled us to test whether the contents of our thoughts could be predicted from brain activation patterns before participants felt they had formed the actual thought, and if the thought content shared characteristics with perceptual content. We used multi-voxel pattern analysis (MVPA) to decode information contained in spatial patterns of brain activation recorded using fMRI (*4*–*6*).

We found that activity patterns were predictive of thought content as far back as 11 seconds before the conscious decision of what to think of –in visual, frontal and subcortical areas. Importantly, predictive patterns in visual and frontal areas contained similar information (i.e., consistent spatial patterns of activation) to representations elicited by unattended sensory stimulation.

Our paradigm consisted of a mental decision leading to the formation of a visual thought or mental image. In every trial, participants had to choose to imagine one of two possible different colored and oriented gratings while we recorded brain blood-oxygen-level dependent (BOLD) using fMRI (Figure 1, see Supplementary Methods for details). After the start of the trial, participants had a maximum of 20 seconds to freely decide which pattern to think of. As soon as they felt they had made the decision, they pressed a button (always the same button for both gratings) with the right hand, thus starting 10 seconds of imagery generation. During this time, participants imagined the chosen grating as vividly as they could. Subsequently, they were prompted with two questions: “what did you imagine?” and “how vivid was it” to which they answered by pressing different buttons (Figure 1). On average, participants took 5.48s (±0.15 SEM) to decide which grating to imagine, while the average trial time was 31.18s (see Figure S1 and Supplementary Methods for details). Each trial included a blank period of 10s at the end to avoid spillover effects from one trial to the next (*7*, *8*). Participants chose to imagine each grating with similar probabilities (50.44% versus 49.56% for vertical and horizontal respectively, see Supplementary Methods for detailed behavioral results).

**Figure 1.**
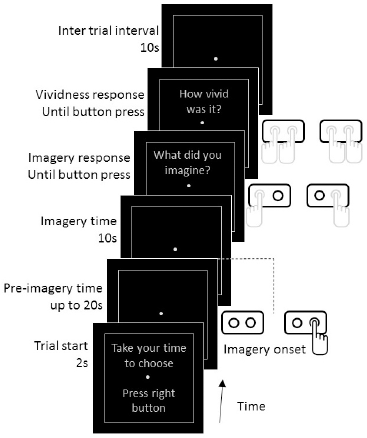
fMRI task paradigm. Participants had to freely choose between two predefined gratings (horizontal green/vertical red or vertical green/horizontal red, counterbalanced across participants). Each trial started with the prompt: “take your time to choose – press right button” for 2 seconds. While the decision was made, a screen containing a fixation point inside a rectangle was shown. This period is referred as “pre-imagery time” and was limited to 20 seconds. Participants were instructed to press a button on the right hand as soon they decided which grating to imagine (always the same button independently of the chosen grating). During the imagery period (10 seconds), participants imagined the chosen grating as vividly as possible. At the end of the imagery period, a question appeared on the screen: “what did you imagine? – Left for vertical green/red – Right for horizontal red/green” (depending on the pre-assigned gratings for the participant). After giving the answer, a second question appeared: “how vivid was it? – 1 (low) to 4 (high)”. After each trial there was a blank interval of 10 seconds where we instructed the participants to relax and not to think about the gratings nor subsequent decisions. Gray hand drawings represent multiple possible button responses, while black drawing represents a unique button choice.

We first verified the suitability of our decoding approach to classify the contents of visual perception and imagery. We thus used SVM trained and tested (in a cross-validation scheme) on 10s of perception or imagery data and classified the perceptual or imagined stimuli (red/green horizontal/vertical gratings) on visual areas from V1 to V4 (see Supplementary Methods for details). Figure S2 shows the results of this sanity check. We found above chance decoding accuracy for perception (91.7, 91.7, 91.7 and 71.4%; one-tailed t-test p = 3.1·10^−8^, 1.2·10^−9^, 7·10^−11^ and 1.5·10^−3^; from V1 to V4) and imagery (66.9, 67, 69.1 and 63.7%; p = 8·10^−4^, 1.2·10^−3^, 1·10^−4^ and 8·10^−3^). These results are comparable to previous results on decoding perception and imagery (*9*–*11*) and thus validate our decoding approach.

To investigate which brain areas contained information about the contents of imagery, we employed a searchlight decoding analysis on the fMRI data (*6*). We used two sources of information to decode the contents of imagery: brain activation patterns from imagery and patterns from perceptual stimuli without attention. For the imagery decoding, we trained classifiers using the imagery data; while in the imagery-perception generalization analysis we trained the classifiers using data from the perception scans and tested on imagery. This allowed us to explore shared information between perception and imagery, without the effects of attention (see Supplementary Methods for details & behavioral attention task during perception).

We defined the areas bearing information about the contents of imagery as those revealing above chance decoding accuracy at any point in time during a trial, thus preventing biasing the results towards the pre-imagery period (cluster definition threshold p<0.001, cluster threshold p<0.05, see Supplementary Methods for details). We thus found a network of four areas: frontal, occipital, thalamus and pons (Figure 2, central panels, see Table S1 for cluster locations in MNI coordinates). We then examined the time course of these areas from −13 to +13 seconds from the reported imagery decision (time = 0). As expected, time-resolved (2s) decoding yielded lower (but statistically significant) accuracies than averaging over longer periods (see Figure S2 for comparison), presumably due to its lower signal-to-noise ratio. Importantly, in the context of neuroscience research, decoding accuracy scores are not the most relevant output of classification, but rather their statistical significance is (*12*). Time-resolved classification in the imagery condition reached above chance decoding accuracy up to 11 seconds before reported imagery onset in occipital, frontal, thalamus and −9 seconds in the pons (Fig. 2; black solid points, p<0.05, one-sample, one-tailed t-test). The perception-imagery generalization decoding showed significant above chance accuracy as early −9 seconds before the onset of imagery in occipital areas and −3 seconds in frontal areas, indicating that pre-volitional predictive information shares properties with perceptual representations in these cortical areas (Fig. 2; grey solid points). In subcortical areas, above-chance generalization decoding accuracy was only observed after the onset of imagery (+1 and +11 seconds in the thalamus and the pons respectively). Importantly, during the perceptual scans visual attention was diverted by a demanding fixation task (see Supplementary Methods), hence such generalization should not be due to high-level volitional or attentional mechanisms. Interestingly, decoding accuracy in occipital areas during the imagery period was lower than expected (see for example (*13*)). Previous studies have shown that prior decisions can impair subsequent cognitive tasks (*14*). Therefore, the cognitive load for the decision element of our task could impair imagery, which is consistent with the results of a behavioral control experiment showing that cued imagery (no-decision) was stronger than decision+imagery (Figure S3B-C).

**Figure 2.**
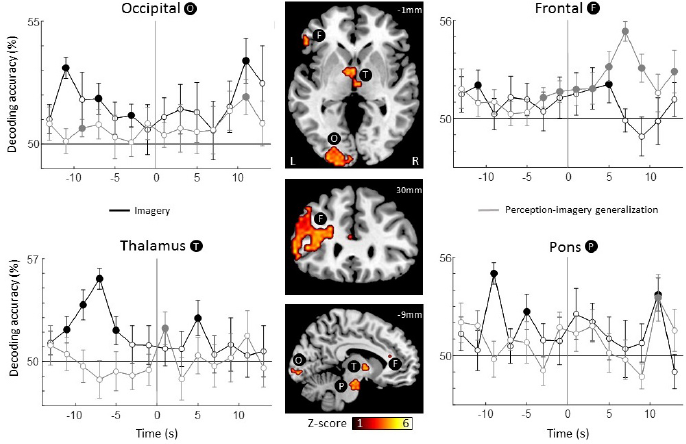
Searchlight decoding of the contents of imagery. Using searchlight decoding we investigated which regions contained information about the contents of mental imagery (see Supplementary Methods for details). We defined these regions as those showing above chance accuracy at any point in time (Gaussian random field correction for multiple comparisons, see Supplementary Methods for details). We found 4 such regions (central panels): occipital (O), frontal (F), thalamus (T) and pons (P). Then, we investigated the temporal dynamics of each one of these regions (lateral plots), from −13 to +13 seconds from imagery onset (time = 0). We decoded imagery contents using the information from imagery runs (imagery, black line) and using information from perception (perception-imagery generalization, grey line). For the imagery decoding (black), all four regions showed significant above-chance accuracy both before and after imagery onset, indicating that information from imagery was predictive of the chosen grating before (up to −11 seconds) and after the imagery onset. On the other hand, the perception-imagery generalization (grey) showed significant above-chance decoding before the onset of imagery only in occipital and frontal areas, indicating that perceptual-like information was predictive of the chosen grating before the imagery onset only in cortical areas (occipital and frontal) and after the imagery onset in both cortical and subcortical areas. Numbers on upper-right slices’ corners indicate MNI coordinates. Error bars represent SEM across participants. Full points represent above chance decoding (p<0.05, one-tailed t-test).

To control for the multi testing problem across time points, we estimated the probability of obtaining above chance significant decoding on any number of time points by chance using permutation tests (see Figure S4 and Supplementary Methods for details). For the imagery condition, the number of decoding time-points was significantly above chance for occipital, thalamus and pons (p<0.05, permutation test, see Figure S4 for details). For the generalization condition, the number of such time-points in frontal was above-chance. Thus, predictive information could be detected confidently with searchlight decoding in occipital, and subcortical areas; while perception-like representations were found in frontal areas.

We conducted a series of controls to test the reliability of our results. First, we ran an independent behavioral experiment to test whether participants might have begun imagining before they reported having done so, which could explain early above chance classification. Outside the scanner, we utilized a method that exploits binocular rivalry to objectively measure imagery strength (15, 16) as a function of time in a free decision and a cued condition (i.e., imposing which stimuli to imagine and when, Figure S3A, See Supplementary Methods for details). We reasoned that if participants were reporting the onset of imagery a few seconds late, this would be detected as an increase in rivalry ‘priming’ compared to a condition where the onset of imagery is imposed, as such priming is known to be dependent on time (*15*). We found the inverse pattern: priming in the free decision condition was significantly lower than in the cued condition (ANOVA, F = 5.77, p = 0.021, Figure S3B), thus ruling out that participants started imagining before they reported imagery onset, and also potentially suggesting that sensory priming might be disrupted by the free decision task, perhaps due to increased cognitive load. Subjective imagery vividness showed a similar trend (Figure S3C, see Supplementary Methods for details). Importantly, we found that this behavioral task has a maximal temporal resolution comparable to that of fMRI (i.e., about 3 seconds). This control largely overcomes one of the limitations to prior free-choice paradigms, as it enables us to measure precision of thought-choice reporting (*17*).

Further we employed a permutation test to check whether the decoding distributions contained any bias, in which case above chance decoding would be overestimated and the use of standard parametric statistical tests would be invalid (*18*) (see Supplementary Methods for details). Permutation tests yielded similar results to those using parametric tests (Figure S5), and, importantly, decoding accuracy distributions under the null hypothesis showed no bias, thus validating the use of standard parametric statistical tests (Table S2). We also conducted a control analysis to test whether the searchlight results could be explained by any spill over from the previous trial. We trained the classifiers on the previous trial (N-1 training) and tested on the subsequent trial (trial N). If there was spill over from the previous trial, this analysis should show similar or higher decoding accuracy in the pre-imagery period. We found no significant above chance classification for any of the regions, thus ruling out the possibility that these results are explained by any spill over (Figure S6).

To investigate how the different regions (occipital, frontal, thalamus and pons) found in the searchlight analysis interact with each other, we used dynamic causal modelling (DCM). We tested whether the source of imagery switched from occipital to frontal during the pre-imagery and imagery periods, and, if the connectivity schemes changed. Specifically, whether occipital and subcortical coupling changes along with the connectivity between frontal and occipital. We thus tested 4 different candidate connectivity schemes (Figure S7.A, see Supplementary Methods for details). Data from before the decision was best explained by an occipital source of imagery, with feedforward connections to frontal and reciprocal connections between the visual and subcortical nodes (Model 4, Figure S7.A). In contrast, data from after the decision was best explained by a frontal source of imagery and occipital-subcortical disconnection scheme (Model 1, Figure S7.A & B).

We then explored the dynamics of the contents of visual imagery along the visual hierarchy. To this purpose, we employed a region-of-interest (ROI) analysis focusing on the first 4 visual cortical areas (V1, V2, V3 and V4) which were identified using standard functional retinotopic mapping (see Supplementary Methods for details). This analysis revealed similar temporal dynamics to the searchlight approach, showing earliest above-chance decoding accuracy −11 seconds from the reported imagery decision (Figure 3A). We found no evident hierarchical pattern in the involvement of visual areas as a function of time (i.e., lower visual areas recruited earlier than higher ones). In the imagery decoding, all visual ROIs showed above chance decoding accuracy before the imagery onset at different time points, however V1 was consistent across time points (from −11 to −5 seconds). The perception-imagery generalization showed more modest effects with above chance decoding accuracy consistently found in V3 as early as 3s before imagery onset (Figure 3B). Permutation tests on the ROI decoding yielded similar results (Figure S8). We controlled for whether these results could be accounted by any spill over from previous trials by again conducting an N-1 analysis. This analysis did not show any above chance accuracy before the imagery onset, but we found a significant time point at t=5s in V4 for the imagery condition (Figure S9). We estimated the family-wise error rate across time points. For the imagery decoding, the number of above-chance decoding time points was significantly above chance for V1 and V4 (p<0.05, permutation test, see Figure S4, for details); while for the generalization only V3 was significantly above chance (p<0.05, Figure S4).

**Figure 3.**
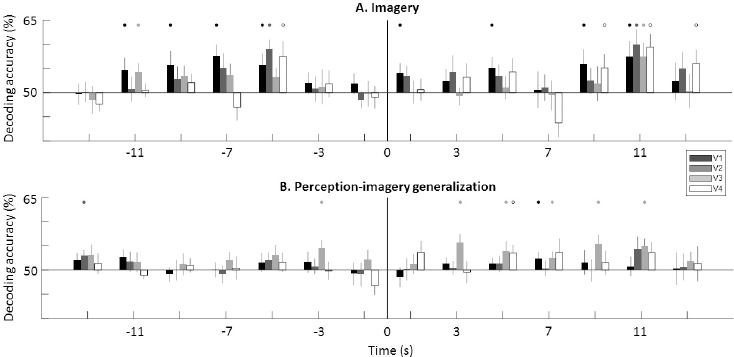
Decoding the contents of imagery in visual regions-of-interest (ROI). We examined the contents of imagery in visual areas using a ROI approach. Visual areas from V1 to V4 were functionally defined and restricted to the foveal representation (see Supplementary Methods for details). **A**. We found above-chance decoding accuracy for imagery decoding both before (from −11 seconds) and after imagery onset. Different visual ROI showed significant above-chance decoding accuracy at different time points, while V1 ROI was the most consistent across time points. **B**. The perception-imagery generalization showed consistent above chance decoding accuracy in V3. Error bars represent SEM across participants. Full points represent above chance decoding (p<0.05, one-tailed t-test).

Next we investigated the effect of subjective imagery vividness on decoding accuracy for imagery. We divided the trials into low-and high-vividness (mean split, see Supplementary Methods for details). As expected, the decoding accuracy for imagery content was higher in high-vividness trials, but surprisingly the biggest differences were observed before the onset of imagery (Figure S10.A). The generalization analysis showed the same trend. We found above chance decoding only in high vividness trials (Figure S10.B), suggesting that in more vivid imagery trials, shared representations between perception and imagery emerge before volition. This result suggests that nonconscious sensory representations in visual areas are present before the onset of conscious thoughts and also that they affect the strength of subsequent conscious imagery.

If the prior, pre-imagery sensory representations in early visual cortex do indeed dictate the strength of subsequent thoughts, then the pre-imagery data should predict the reported vividness from the subsequent imagery period. Accordingly, we tested exactly this, we attempted to decode the subjective strength of the imagery (i.e., vividness) by using only the fMRI data from before the imagery period. Decoding accuracy was significantly above chance in V1 (62.2%, p=0.0035, one-tailed t-test), but not in other visual ROIs (p>0.05, Figure 4), indicating that information contained in V1 predicted future subjective imagery strength.

**Figure 4.**
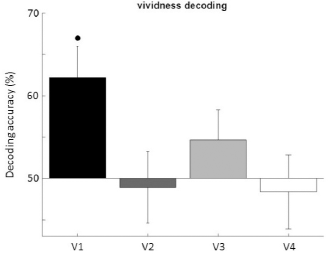
Pre-imagery brain activity predicts the strength of subsequent visual thoughts. We used pre-imagery data to decode subsequent imagery vividness (see text for details). Information in V1 from before the imagery decision predicted how vivid the future imagery will be. Error bars represent SEM across participants. Full point represents above chance decoding (p<0.05, one-tailed t-test).

To summarize, we found that brain patterns were predictive of the contents of imagery as far as -11 seconds before the choice of what to imagine. These results indicate that the contents of future thoughts can be biased by nonconscious representations. Our results show predictive patterns in occipital, frontal and subcortical areas. While previous results highlight the role of frontal areas carrying information about subsequent decisions (*19*–*21*); to the best of our knowledge, visual and subcortical areas have not been reported. Interestingly, here we found that predictive information in visual and frontal areas shared similarities with representations elicited by sensory stimulation, suggesting that perception-like neural representations are present before the decision of what to imagine. In previous experiments applying MVPA to study decision processes, predictive information about choices has been interpreted as evidence for nonconscious decision making (*19*, *20*, *22*). Thus, our results could be interpreted as the imagery decision being made (at least partly) nonconsciously, supporting the idea that subjective sensation of making the decision emerges after the decision is already made (*19*, *20*, *23*, *24*). However, an alternative hypothesis is that these results reflect decisional mechanisms that rely on spontaneously generated subliminal visual representations present before the decision. Since the goal of the task was to randomly choose and imagine a grating as vividly as possible, one strategy might be to choose the pattern that is already more strongly represented, albeit subliminally. In other words, spontaneous grating representations might stochastically fluctuate in strength while remaining weak enough to escape awareness. Thus, prior to the decision, one grating representation might dominate, hence being more prone to decisional thought-selection. A similar interpretation has been advanced to explain the buildup of neural activity prior to self-initiated movements (aka readiness potential) (*25*). Interestingly, it has been recently shown that self-initiated movements can be aborted even after the onset of predictive neural signals (*26*), suggesting that the decision can be somewhat dissociated from predictive neural signals. Therefore, our results can be explained by a conscious choice that relies on nonconscious representations during the decision production; perhaps analogous to blindsight (*27*), subliminal priming studies (28, 29) or nonconscious decisional accumulation (*30*). Such a mechanism is intriguing in light of theories of mental imagery and thought generation that propose involuntary thought intrusion as both an everyday event, and, in extreme cases, a component of mental disorders like PTSD (31, 32).

We found support for this hypothesis in our exploratory DCM analysis (Figure S7, see Supplementary Methods for details). Effective connectivity analysis results suggest, that during the pre-imagery period, non-conscious visual representations in occipital areas are read by frontal areas; while occipital cortex is coupled with subcortical regions, perhaps triggering the stochastic fluctuations in grating representation strength. This is consistent with the finding that the thalamus is functionally coupled with the visual cortex, especially in resting state mode (33, 34). While, anatomical connections between the pons and visual areas have been found in primates (*35*), its functional role is still unclear. On the other hand, during the subsequent imagery period, occipital and subcortical areas would be uncoupled, thus stabilizing the contents of imagery, volitionally led by frontal areas (Figure S7.B).

Accordingly, our current study can be seen as the first to capture the possible origins and contents of involuntary thoughts and how they progress into or bias subsequent voluntary conscious thoughts. This is compatible with the finding that the most prominent differences between low and high vividness trials are seen for the pre-imagery period in visual areas, which can be interpreted as when one of the patterns is more strongly represented (subliminally) it will induce a more vivid subsequent volitional image.

It is up to future research to dissociate between the two hypotheses of nonconscious decisions and nonconscious prior representations biasing subsequent fully conscious volitional decisions. Untangling these models will not only shed light on age-old questions of free will, but also provide a clear mechanism for pathological intrusive thoughts common across multiple mental disorders.

## Acknowledgements

We would like to thank Johanna Bergmann for her input in the experimental design, useful comments and help with participant’s testing. Eugene Kwok for his help in the behavioral testing. Collin Clifford, Damien Mannion and Kiley Seymour for useful comments. This research was supported by Australian NHMRC grants GNT1046198 and GNT1085404 and JP was supported by a Career Development Fellowship GNT1049596 and ARC discovery projects DP140101560 and DP160103299.

## Supplementary methods

### Participants

Experimental procedures were approved by the University of New South Wales Human Research Ethics Committee (HREC#: HC12030). All participants gave written consent to participate in the experiment. For the behavioral free decision and cued imagery priming task, we tested 8 participants (4 females, aged 29.3±0.5 years old). For the fMRI experiment, we tested 14 participants (9 females, aged 29.1±1.1 years old).

### Behavioral imagery decision precision control experiment

Since self-report of the onset of decision has been criticized due to its unreliability (*1*), we developed an independent psychophysics experiment to test its reliability. We objectively measured imagery strength as a function of time for a subset of the participants of the fMRI experiment. We employed two conditions: free decision (freely chosen imagined stimulus and imagery onset) and cued (i.e., imposed imagined stimuli and imagery onset). We used binocular rivalry priming as a means to objectively measure sensory imagery strength (*2*–*4*). When imagining one of the competing stimuli prior to binocular rivalry, the rivalry perception is biased towards the imagined stimulus, with a stronger priming as the imagery time increases (*2*); see (*2*, *5*) for discussion of why this is an objective measure. We asked participants to imagine one of the gratings for different durations and then measured rivalry priming as a function of the different durations. We reasoned that if participants reported the onset of imagery a few seconds after they actually started imagining, this would be detected as an increase in priming compared to the condition where the onset of imagery is imposed. Thus, in the free decision condition, participants had to freely choose to imagine one of the two predefined gratings (horizontal green/vertical red or vertical green/horizontal red, counterbalanced across participants). In the cued condition, participants were presented with a cue indicating which grating to imagine, thus imposing the onset of imagery as well as which grating needed to be imagined. Each trial started with the instruction “press space bar to start the trial” (Figure S3A). Then, either the instruction “CHOOSE” or a cue indicating which grating to imagine (i.e., “horizontal red” was presented for 1 second. In the free decision condition, the imagery time started after the participant chose the grating to imagine, which was indicated by pressing a key on the computer keyboard (Figure S3A). For the cued imagery, the imagery time started right after the cue was gone (i.e., no decision time). We tested 3 imagery times (3.33, 6.67 and 10 seconds). After the imagery time, a high pitch sound was delivered (200ms) and both gratings were presented through red/green stereo glasses at fixation for 700ms. Then, participants had to report which grating was dominant (i.e., horizontal red, vertical green or mixed if no grating was dominant), by pressing different keys. After this, they had to answer which grating they imagined (for both free decision and cued trials). Participants then rated their imagery vividness from 1 (low) to 4 (high). Free decision and cued trials as well as imagery times were pseudo-randomized within a block of 30 trials. We added catch trials (20%) in which the gratings were physically fused and equally dominant to control the reliability of self-report (2, 6). We tested 120 trials for each free decision and cued imagery conditions (40 trials per time point), plus 48 catch trials evenly divided among time points. Priming and vividness were normalized as z-score within participants and across time-points and conditions, to account for baseline differences across participants, but otherwise conserving relative differences amongst conditions and time-points.

Figure S3B shows the effects of imagery time on sensory priming for both conditions. Imagery time showed a significant effect on priming for free decision and cued conditions (ANOVA, F = 7.15, p = 0.002, Figure S3B), thus confirming the effect of imagery time on priming. Priming for the free decision condition was significantly lower than in the cued condition (ANOVA, F = 5.77, p = 0.021), indicating that participants were not starting imagining before they reported doing so (which would have resulted in the opposite pattern) and also suggesting that sensory priming is somehow disrupted by the decision task, perhaps due to cognitive load. Importantly, significant differences in priming between 3.33 and 6.67 seconds of imagery time were found for the free decision and cued conditions (one-tailed t-test, p<0.05), indicating that this behavioral task can resolve differences in priming spaced by 3.33 seconds, at least for these two first time points, thus providing a lower bound of temporal resolution of the accuracy of the reported imagery onset. For further information on why this method measures imagery strength and not visual attention, binocular rivalry control or response bias see (*2*, *5*). Self-report reliability was verified with catch trials which were reported as mixed above chance level (83.8%, p=0.002).

Figure S3C shows the effects of imagery time on subjective imagery vividness. Imagery time showed also a significant effect on vividness for free decision and cued conditions (ANOVA, F = 18.49, p < 10^-5^, Figure S3C). However, differences between free decision and cued conditions were not significant (ANOVA, F = 2.42, p = 0.127). Again, significant differences in vividness between 3.33 and 6.67 seconds of imagery time were found for the free decision and cued conditions (one-tailed t-test, p < 0.01), setting thus a lower bound of 3.33 seconds for the temporal resolution on the behavioral task.

We tested this independent behavioral experiment on 8 randomly chosen participants from the fMRI experiment, who had extensive experience as subjects in psychophysics experiments. We sought to test if these results would generalize to completely unexperienced participants in psychophysics experiments (N=10). We did not, however, find significant results (not shown), suggesting that this is a highly demanding task and experience in psychophysics might be important to perform the task properly.

Functional and structural MRI parameters. Scans were performed at the Neuroscience Research Australia (NeuRA) facility, Sydney, Australia, in a Philips 3T Achieva TX MRI scanner using a 32-channel head coil. Structural images were acquired using turbo field echo (TFE) sequence consisting in 256 T1-weighted sagittal slices covering the whole brain (flip angle 8 deg, matrix size = 256×256, voxel size = 1mm isotropic). Functional T2*-weighted images were acquired using echo planar imaging (EPI) sequence with 31 slices (flip angle = 90 deg, matrix size = 240x240, voxel size = 3mm isotropic, TR = 2000ms, TE = 40ms).

### fMRI free decision visual imagery task

We instructed participants to choose between two predefined gratings (horizontal green/vertical red or vertical green/horizontal red, counterbalanced across participants), which were previously familiar to the participants, through prior training sessions. We asked the participants to refrain from following preconceived decision schemes. In the scanner, participants were provided with two dual-button boxes, one held in each hand. Each trial started with a prompt reading: “take your time to choose – press right button” for 2 seconds (Figure 1). After this, a screen containing a fixation point was shown while the decision as to what to think of was made. This period is referred as “pre-imagery time” and was limited to 20 seconds. Participants were instructed to press a button with the right hand as soon as they decided which grating to imagine. After pressing the button, the fixation point became brighter for 100ms to indicate the participants that the imagery onset time was recorded. During the imagery period (10 seconds), participants were instructed to imagine the chosen pattern as vividly as possible trying, if possible, to project it onto the screen. At the end of the imagery period, a question appeared on the screen: “what did you imagine? – Left for vertical green/red – Right for horizontal red/green” (depending on the pre-assigned patterns for the participant). After giving the answer, a second question appeared: “how vivid was it? – 1 (low) to 4 (high)”. After each trial there was a blank interval of 10 seconds where we instructed the participants to just relax and try not to think about the gratings nor any subsequent decisions. Post-experiment interviews revealed that some participants (n=4) could not help thinking about gratings in some trials during the inter trial interval. They reported different strategies to avoid these thoughts such as ignoring them, replacing them for another image/thought, or choosing the other grating when the decision came. The remaining participants (n=10) reported not having any thoughts or mental images about gratings during the rest period. We tested if the effects we found could be explained by the former group of participants who could not refrain from thinking about gratings. We thus performed the analysis using only data from the participants who did not think/imagine gratings outside the imagery period (n=10). Figure S12 shows the results of this control. Results are comparable to those shown in Figure 2, thus ruling out the possibility that that the effects we report were driven by the 4 participants who had troubles dis-engaging from imagery in the rest period. We delivered the task in runs of 5 minutes during which the participants completed as many trials as possible. Participants chose to imagine horizontal and vertical grids with a similar probability (50.44% versus 49.56% for vertical and horizontal gratings respectively, mean Shannon entropy = 0.997±0.001 SEM) and showed an average probability of switching gratings from one trial to the next of 58.59% ±2.81 SEM.

### fMRI perception condition

We presented counter-phase flickering gratings at 4.167 Hz (70% contrast, ~0.5 degrees of visual angle per cycle). They were presented at their respective predefined colors and orientations (horizontal green/vertical red or vertical green/horizontal red). The gratings were convolved with a Gaussian-like 2D kernel to obtain smooth-edged circular gratings. Gratings were presented inside a rectangle (the same that was used in the imagery task, Figure 1) and a fixation point was drawn at the center (as for the imagery task). Within a run of 3 minutes, we presented the flickering patterns in a block manner, interleaved with fixation periods (15 seconds each). Importantly, an attention task was performed consisting of detecting a change in fixation point brightness (+70% for 200ms). Fixation changes were allocated randomly during a run, from 1 to 4 instances. Participants were instructed to press any of the 4 buttons as soon as they detected the changes. Participants showed high performance in the detection task (d-prime=3.33 ±0.13 SEM).

### Functional mapping of retinotopic visual areas

To functionally determine the boundaries of visual areas from V1 to V4 independently for each participant, we used the phase-encoding method (*7*, *8*). Double wedges containing dynamic colored patterns cycled through 10 rotations in 10min (retinotopic stimulation frequency = 0.033 Hz). To ensure deployment of attention to the stimulus during the mapping, participants performed a detection task: pressing a button upon seeing a gray dot anywhere on the wedges. The script for this experiment was downloaded from Samuel Schwarzkopf’s tutorial (http://sampendu.wordpress.com/retinotopy-tutorial/").

### Experimental procedures

We performed the 3 experiments in a single scanning session lasting about 1.5h. Stimuli were delivered using an 18” MRI-compatible LCD screen (Philips ERD-2, 60Hz refresh rate) located at the end of the bore. All stimuli were delivered and responses gathered employing the Psychtoolbox 3 (*9*, *10*) for MATLAB (The MathWorks Inc., Natick, MA, USA) using inhouse scripts. Participants’ heads were restrained using foam pads and adhesive tape. Each session followed the same structure: first the structural scanning followed by the retinotopic mapping. Then the perception task was alternated with the imagery task until completing 3 runs of the perception task. Then the imagery task was repeated until completing 7 or 8 (depending on the participant) runs in total. Pauses were assigned in between the runs. The 4 first volumes of each functional runs were discarded to account for the equilibrium magnetization time and each functional run started with 10 seconds of fixation.

### Phase-encoded retinotopic mapping analysis

fMRI retinotopic mapping data were analyzed using the Fast-Fourier Transform (FFT) in MATLAB. The FFT was applied voxel-wise across time points. The complex output of the FFT contained both the amplitude and phase information of sinusoidal components of the BOLD signal. Phase information at the frequency of stimulation (0.033Hz) was then extracted, using its amplitude as threshold (≥2 SNR) and overlaid them on each participant’s cortical surface reconstruction obtained using Freesurfer (*11*, *12*). We manually delineated boundaries between retinotopic areas on the flattened surface around the occipital pole by identifying voxels showing phase reversals in the polar angle map, representing the horizontal and vertical visual meridians. In all participants, we clearly defined five distinct visual areas: V1, V2, V3d, V3v and V4; throughout this paper, we merge V3d and V3v and label them as V3. All four retinotopic labels were then defined as the intersection with the perceptual blocks (grating>fixation, p<0.001, FDR corrected) thus restricting the ROI to the foveal representation of each visual area.

### fMRI signal processing

All data were analyzed using SPM12 (Statistical Parametric Mapping; Wellcome Trust Centre for Neuroimaging, London, UK). We realigned functional images to the first functional volume and high-pass filtered (128 seconds) to remove low-frequency drifts in the signal. To estimate the hemodynamic response function (HRF), we generated regressors for each grating (horizontal green/vertical red or vertical green/horizontal red) for each run and experiment (perception and imagery) independently. We used finite-impulse response (FIR) as the basis function that makes no assumptions about the shape of the HRF which is important for the analysis of the free decision imagery data (*13*). We employed a 14th order FIR basis function encompassing 28 seconds from -13 to +13 seconds from the imagery onset, thus obtaining 14 bins representing each TR. For the perception condition, we employed a 1^st^ order FIR basis function from the onset of each perceptual block to its end (15 seconds). For the vividness analysis, we split the trials into low-vividness (ratings 1 and 2) and high-vividness (ratings 3 and 4), we then obtained the regressors for both gratings as explained above.

### Multi-voxel pattern analysis (MVPA)

We used a well stablished decoding approach to extract information related to each grating contained in the pattern of activation across voxels of a given participant using the decoding toolbox (TDT) (*14*). Using a leave one-run out cross-validation scheme, we trained a linear supporting vector machine (SVM) on all runs but one and then tested on the remaining one. We repeated this procedure until all runs were used as test and then averaged the results across validations (7 or 8-fold, depending on the participant). We performed one-run leave out cross validation for every temporal bin independently. We also employed cross-classification to generalize information between the perception and the imagery tasks. For the cross-classification, we trained on the ensemble of the perception runs and tested on the ensemble of the imagery runs. We employed 2 different decoding approaches: searchlight and region-of-interest (ROI). We used a spherical searchlight of 3 voxels of radius and obtained volumes in which a value of decoding accuracy was assigned to each voxel. We normalized the decoding accuracy volumes into the MNI space and applied a spatial smoothing of 8mm FWHM. We then performed a one-tail one-sample t-test against 50% (chance level) across participants for every voxel. We corrected for multiple comparisons using cluster-extent based thresholding employing Gaussian Random Field theory (*15*, *16*), as implemented in FSL (*17*). We used a conservative primary threshold of p<0.001 at the voxel level, as recommended in previous studies (*18*), and a cluster level threshold of p<0.05 in every time point volume independently. Importantly, these thresholds have been shown to be valid within the nominal false positive ratios (*19*). ROI decoding was used to test information content in visual areas specifically. We defined the boundaries of visual areas from V1 to V4 which volumes were used as ROI. Note that because visual ROI were defined on the cortical surface (see phase-encoded retinotopic analysis for details), only gray-matter containing voxels were considered, as opposed to the searchlight approach which also considers non-gray matter containing voxels. We tested if there was a difference in the average BOLD response between stimuli (i.e., univariate difference). We did not find any significant differences (p>0.05, uncorrected) in the average BOLD response (Figure S11), thus ruling-out the possibility that the results would be explained by differences in the average level of activity across conditions.

### Permutation test

In order to validate the use of standard parametric statistics we performed a permutation test and thus empirically determined the distribution of decoding accuracies under the null hypothesis (*20*). Previous reports have highlighted the possibility of obtaining skewed decoding distributions, which would invalidate the use of standard parametric statistical tests (*21*). We thus randomly shuffled the labels (i.e., horizontal red/vertical green) among trials and within blocks (i.e., number of red/green imagined trials was conserved within a run but trial labels were shuffled) for each participant and condition (imagery and generalization) to generate empirical data under the null hypothesis. After reshuffling the labels, we generated regressors for each stimuli and performed decoding following the same procedure described in the previous paragraph. We repeated this procedure 1000 times and obtained the empirical distribution under the null hypothesis, from which we obtained confidence intervals for each decoding time point and area (Figure S5 and S8) using the percentile method (20). Our results show that the decoding null hypothesis followed a normal distribution (Table S2) and importantly, significant results using permutation test confidence intervals were comparable to the results using standard parametric tests (compare significant points on figures 2 and 3 with figures S5 and S8). This analysis thus validates the use of standard statistical tests to test significance on our dataset.

### Across time-points family-wise error estimation

We estimated the probability of obtaining an n number of significantly above-chance decoding time points (p<0.05, one tailed t-test) under the null hypothesis. To do this, we employed the data from the null distribution obtained with the permutation test (randomly shuffled labels, 1000 iterations; see previous paragraph for details). Figure S4 shows the result of such analysis. Insets show the family-wise error rate for the empirically observed number above-chance decoding time points for each area.

### Dynamic causal modelling (DCM)

We conducted a dynamic causal modelling (DCM) analysis to obtain a putative effective connectivity architecture explaining our results (*22*) (Figure S7). We used 2 variables to construct the candidate models: (1) imagery source and (2) connectivity among areas. For the imagery source, we tested 2 possibilities: frontal source (models 1 and 2) and occipital source (models 3 and 4). As for the connections, we tested either reciprocal occipital-frontal connection with frontal feedback to subcortical areas (models 1 and 3) or reciprocal occipital-subcortical connections and feedforward from occipital to frontal. We chose these connectivity schemes to test a simple hypothesis: whether occipital and subcortical nodes were coupled in the pre-imagery period, which would explain the stochastic fluctuations in the grating representations strength. And whether this coupling was interrupted by the engagement of frontal areas in the imagery period. We feed independent models with data from before the decision (from -10 to 0 seconds) and after the decision (from 0 to 10 seconds), pooling regressors from red and green gratings. We fitted the models assuming that the model’s structure is conserved across participants, thus employing fixed effects inferences (23). Data from before the decision was best explained by an occipital source of imagery with reciprocal connections between occipital and subcortical nodes, and feedforward connection from occipital to frontal (Model 4, Figure S7). On the other hand, data from after the decision was best explained by a frontal source of imagery and occipital-subcortical disconnection scheme (Model 1, Figure S7). These results suggest that the flow of information changes between the periods before and after the decision. Note that DCM was originally implemented to test effective connectivity in systems receiving perceptual input, thus its validity at inferring connectivity schemes in absence of visual input is unclear—e.g., nonconscious thoughts, mental imagery (however see (24)). Note that our DCM architectures are undiscerning about the contents of imagery (e.g. vertical green and horizontal red) thus only allowing inferences about the global flow of information, as opposed to the contents of imagery (as resolved by decoding analysis). Due to computational and time limitations we did not perform a comprehensive search of all connectivity schemes, thus other models could be as or more suited to explain our data. All in all, we must be cautious at interpreting the connectivity results and they should be taken as an exploratory analysis that complements the decoding analyses.

### Supplementary Figures and Tables

**Figure S1.**
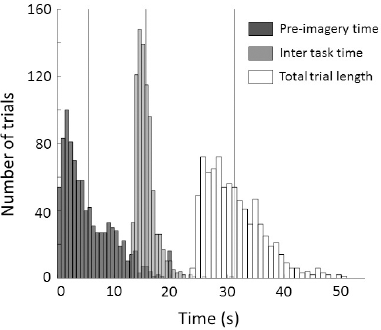
Distributions of the periods composing the free-decision task in the fMRI experiment. Pre-imagery time corresponds to the interval from the end of the trial-start instructions (“take your time to choose / press right button” presented for 2s) to the button press, indicating the start of the imagery period. The Inter task period corresponds to the lapse between the end of the imagery period from one trial to the start of the pre-imagery period in the next trial. Total trial length was defined from onset to onset of the trial instructions in consecutive trials. Vertical lines represent the mean of every period. Average times for pre-imagery, inter task and total trial length were 5.48s, 15.62s and 31.18s, respectively. Note that these time intervals left enough time between trials to avoid activity spill over.

**Figure S2.**
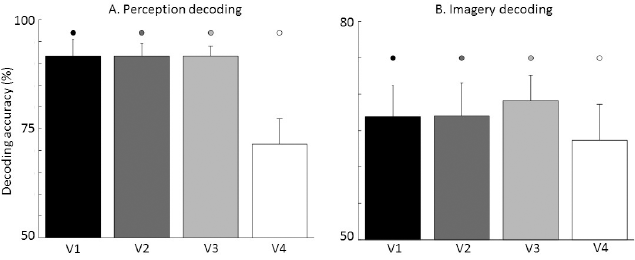
Sanity check of the decoding of perception and imagery contents. We validated our classification approach by decoding perception (A) and imagery (B) on visual ROI. A leave-one-run cross validation scheme was used to train and test linear classifiers (SVM). Perception fMRI data was extracted from 10s of perception blocks. Imagery fMRI data were extracted from the 10s imagery time in the free decision task. Perception (91.7, 91.7, 91.7 and 71.4%; from V1 to V4) and imagery (66.9, 67, 69.1 and 63.7%) decoding accuracy was comparable to previous reports (25–27), thus validating our classification approach. Error bars represent SEM across participants. Dots represent above chance decoding (chance level=50%, p<0.01).

**Figure S3.**
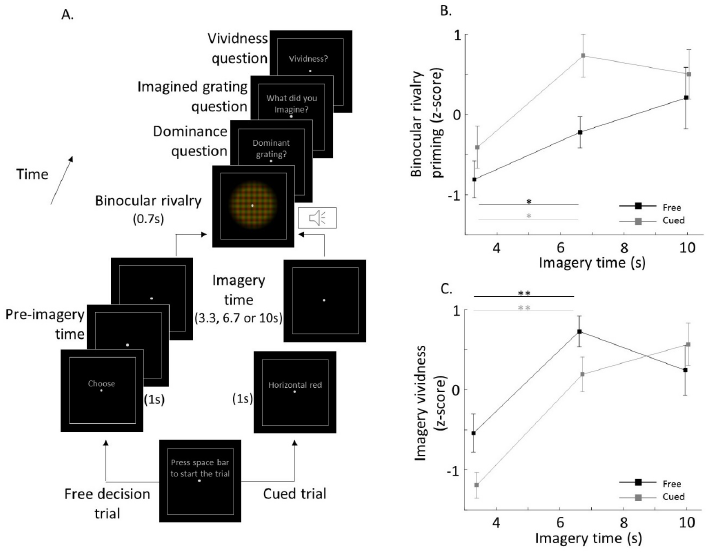
Testing the accuracy of the imagery onset report on a behavioral task. We tested perceptual priming and subjective imagery vividness a function of imagery time as a means to verify the accuracy of reporting the imagery onset. **A. Paradigm**. Free decision and cued trials were pseudo-randomized. Perceptual priming was measured as a function of imagery time (3.3, 6.7 and 10s), as the dominance bias on binocular rivalry. **B. Perceptual priming**. Imagery time significantly increased perceptual priming on the free decision and cued conditions (ANOVA, F = 7.15, p = 0.002), and priming in the free decision condition was significantly lower than in the cued condition (ANOVA, F = 5.77, p = 0.021), thus ruling out that participants were reporting the imagery onset after starting imagining. **C. Imagery vividness**. Imagery time also significantly increased subjective imagery vividness on the free decision and cued conditions (ANOVA, F = 18.49, p < 10^−5^). Stars show significant differences between the first two time points, thus setting a lower bound of temporal resolution on this behavioral task. Error bars show ±SEM.

**Figure S4.**
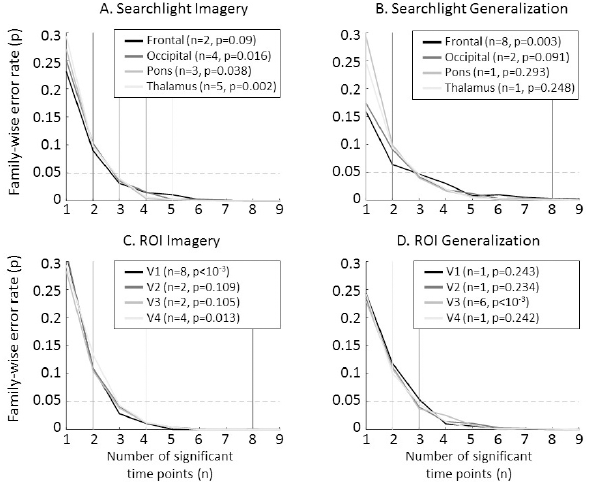
Assessment of the family-wise error rate across time points. We estimated the probability of obtaining different number (n, from 1 to 9) of significantly above chance decoding across time points (p<0.05, one tailed t-test) under the null hypothesis using the null distribution from the permutation test (1000 iterations). Insets show the family-wise error rate for the empirically observed number above-chance decoding time points for each area.

**Figure S5.**
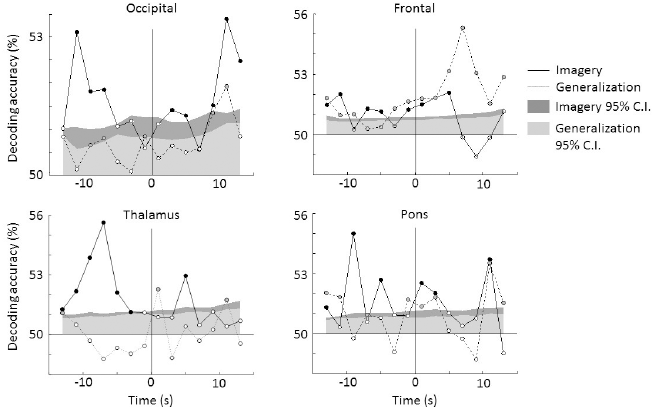
Searchlight confidence intervals (C.I.) from permutation test. We validated the statistical significance of the searchlight decoding accuracy by empirically determining the distribution of the null hypothesis. We thus performed a permutation test (1000 iterations) for each participant and decoding method independently by randomizing the labels (horizontal/vertical, red/green) from trials prior the construction of regressors (see text for details). Confidence intervals (95%, right-tailed) for imagery and generalization are shown in dark and light gray, respectively. Significant above-chance decoding time-points (p<0.05, right-tailed permutation test) are depicted as solid circles. Results are comparable to those shown in Figure 2 in which parametric tests were used (t-test, one-sample, right-tailed), thus validating the use of parametric statistical tests.

**Figure S6.**
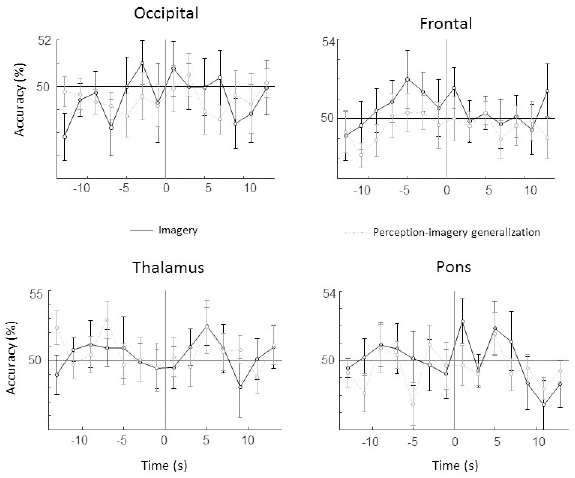
Searchlight spillover effect control. We conducted a control analysis to test whether our results could be explained by activity from the previous trial (spillover effect). We thus trained our classifiers on the previous trials (N-1) and tested on the subsequent one (N). If there was a spill over from previous trial, this analysis should show the same or higher decoding accuracy in the pre-imagery period. We found no significant above chance classification for any of the regions (p>0.05, one-tailed t-test), thus ruling out the possibility that these results are explained by any spill over. Error bars represent ±SEM across participants.

**Figure S7.**
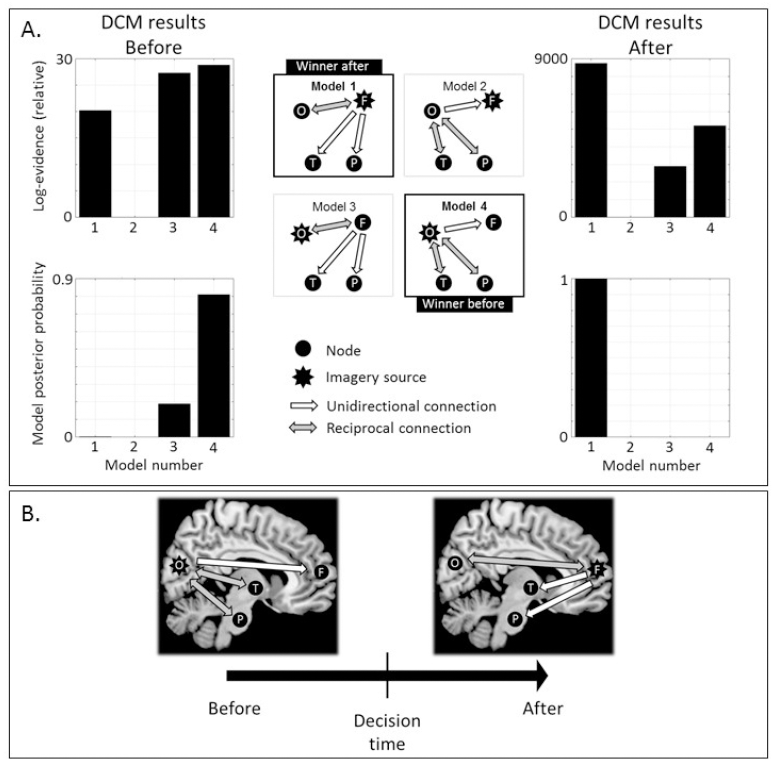
Dynamic causal modelling (DCM) of the source of imagery and its effective connectivity. We used DCM to test different causal architectures accounting for our results. **A**. Clusters from the searchlight analysis were used as nodes: frontal, occipital, thalamus and pons. We tested 2 sources of imagery: frontal, occipital and 2 connectivity schemes. Data from before the decision was best explained by an occipital source of imagery with reciprocal connections between the visual and subcortical nodes (Model 4). Data from after the decision was best explained by a frontal source of imagery and occipital-subcortical disconnection (Model 1). **B**. Summary and functional interpretation. During the pre-imagery period, non-conscious visual representations in occipital areas are read by frontal areas; while occipital is coupled with subcortical regions, perhaps triggering the stochastic fluctuations in gratings representation strength. On the other hand, during the imagery period, occipital and subcortical areas would be uncoupled, thus stabilizing the contents of imagery, which are led by frontal areas.

**Figure S8.**
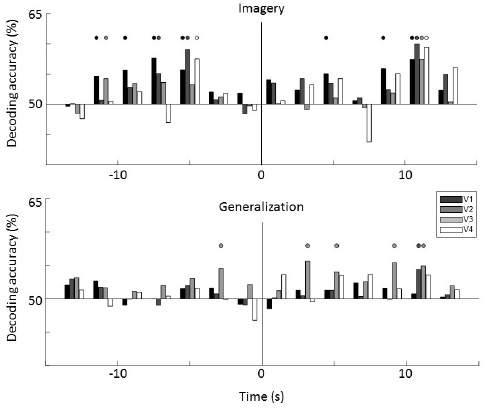
ROI results from permutation test. We validated the statistical significance of the ROI decoding accuracy by empirically determining the distribution of the null hypothesis. We thus conducted a permutation test (1000 iterations) for each participant, decoding method and time-point independently by randomizing the labels (horizontal/vertical, red/green) from trials prior the construction of regressors (see text for details). Significant above-chance decoding time-points (p<0.05, right-tailed permutation test) are depicted as solid circles. Results are comparable to those shown in Figure 3 in which parametric tests were used (t-test, one-sample, right-tailed), thus validating the use of parametric statistical tests.

**Figure S9.**
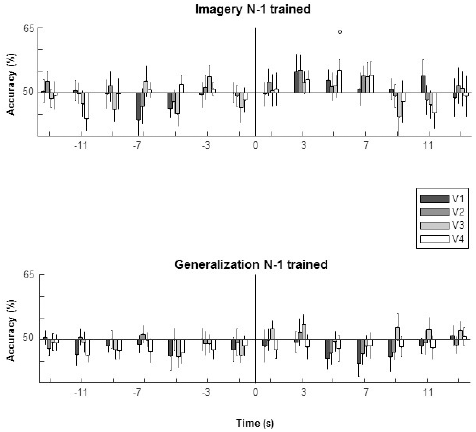
ROI spillover effect control. We conducted a control analysis to test whether our results could be explained by activity from the previous trial (spillover effect). We thus trained our classifiers on the previous trials (N-1) and tested on the subsequent one (N). If there was a spill over from previous trial, this analysis should show the same or higher decoding accuracy in the pre-imagery period. We found no significant above chance classification on the pre-imagery period but we did found significant decoding accuracy in V4 at +5s from imagery onset in the imagery condition. Nevertheless, this result indicates that effects in the pre-imagery time cannot be explained by activity spill over from previous trials. Error bars represent ±SEM across participants.

**Figure S10.**
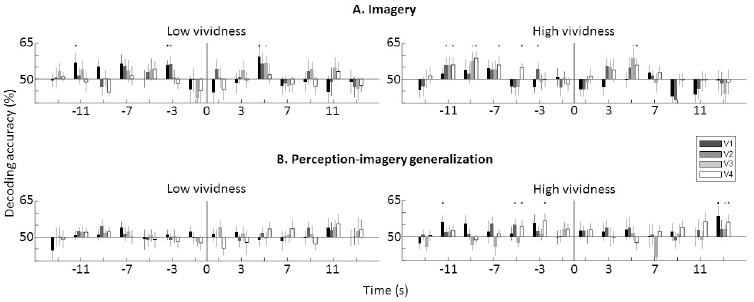
Imagery-content decoding for low-and high-vividness trials. We divided data into trials with low and high vividness (see text for details). High vividness trials showed higher decoding accuracy than low vividness trials. Greatest differences in decoding accuracy as a function of vividness were seen in the pre-imagery period, suggesting that subjective imagery vividness depends upon neural activity from before imagery. Error bars represent SEM across participants. Full points represent above chance decoding (p<0.05, one-tailed t-test).

**Figure S11.**
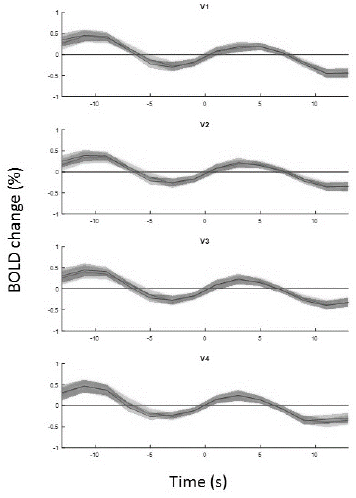
BOLD amplitude change for different imagined gratings. We tested if our effects could be explained by an overall amplitude differences between imagines gratings (i.e., univariate difference). We thus calculated the signal change for every voxel. Dark and light gray represent horizontal and vertical gratings, respectively. BOLD signal changes from both imagined gratings are overlapped and no significant differences were found (p>0.05, t-test, uncorrected). Shade areas represent SEM.

**Figure S12.**
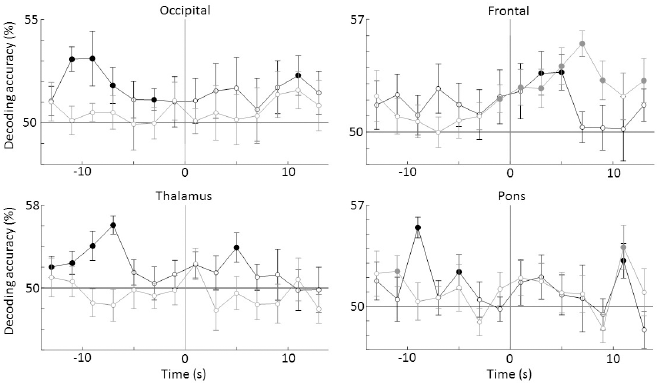
Decoding on a subset of participants. Based in a post-experiment interview we pinpoint some participants (n=4) that could not help thinking about gratings in some trials during the inter trial interval. In order to test if the effects reported here could be explained by these subset of participants, we performed the analysis on the remaining participants (n=10) who reported not having any thoughts or mental images about gratings during the inter trial interval. The control analysis revealed similar results to those presented on Figure 2, thus indicating that our results cannot be explained by the participants who had troubles keeping their minds free from gratings in the inter-trial period. Solid circles represent significant above-chance decoding time-points (p<0.05, right-tailed t-test).

**Table S1.** Searchlight clusters locations. A. Center of mass in MNI coordinates of the 4 clusters found in the searchlight analysis (occipital, frontal, thalamus and pons). B. Structure definitions according to the AAL atlas (*28*), except for the pons label which was defined using Freesurfer’s subcortical automatic segmentation (*29*). Note that Freesurfer’s subcortical segmentation does not define subdivisions within the brainstem. We thus identified the pons by visual inspection of activations within the brainstem, which were mostly located in the brainstem’s anterior protrusion rostral to the medulla, consistent with the pons location (*30*). Percentages represent atlases’ labels volume occupied by the clusters.

|  | A. MNI x,y,z (mm) | B. Atlas labelling |
| --- | --- | --- |
| Occipital | -17,-95,2 | Middle occipital lobe (L) 36.1%<br>Calcarine (L) 27.9%<br>Superior occipital lobe (L) 14.3%<br>Inferior occipital lobe (L) 14.3% |
| Frontal | -36,32,18 | Inferior frontal gyrus, triangular (L) 57.5%<br>Middle frontal gyrus (L) 6.6% |
| Thalamus | 1,-7,5 | Thalamus (R) 27.2%<br>Thalamus (L) 11.2% |
| Pons | 7,-19,-24 | Brainstem 34.8%<br>Cerebellum (R) 9.5%<br>Parahippocampus (R) 5.7% |

**Table S2.** Distribution of the empirically-determined decoding null-hypothesis. We verified the normality of the null-hypothesis decoding distributions (determined using permutation tests, see text for details) by calculating the mean decoding accuracy, skewness and kurtosis. We calculated these values for each cluster in the searchlight analysis: fontal (F), occipital (O), pons (P) and thalamus (T) and each visual ROI: V1, V2, V3 and V4, for the imagery and generalization conditions. The expected decoding accuracy is 50% for the null hypothesis. Expected values of skewness are between -1 and 1; and kurtosis of 3 for normal distributions (31). Our results show that decoding null-hypothesis distributions for both conditions (imagery and generalization) and decoding methods (searchlight and ROI) are centered on 50% and fulfill normal distributions criterion, thus validating the use of standard parametric statistical tests.

|  |  | Searchlight |  |  |  | ROI |  |  |  |
| --- | --- | --- | --- | --- | --- | --- | --- | --- | --- |
|  |  | F | O | P | T | V1 | V2 | V3 | V4 |
| Imagery | Decoding accuracy (%) | 50.0092 | 50.0051 | 49.9879 | 49.9937 | 49.9861 | 49.9982 | 49.9646 | 49.9835 |
|  | Skewness | 0.0658 | 0.0248 | 0.0439 | 0.0243 | -0.05 | -0.033 | -0.0088 | -0.0401 |
|  | Kurtosis | 2.9579 | 2.9974 | 2.9668 | 2.9549 | 2.9106 | 2.9598 | 2.9714 | 2.9503 |
| Generalization | Decoding accuracy (%) | 49.9926 | 49.9985 | 50.0033 | 50.0079 | 50.0203 | 50.0457 | 49.9951 | 49.9533 |
|  | Skewness | -0.0514 | -0.0135 | 0.0039 | -0.0212 | -0.0251 | -0.0162 | -0.0237 | 0.0046 |
|  | Kurtosis | 3.0005 | 2.9663 | 2.9476 | 3.0354 | 2.9661 | 2.9996 | 3.0071 | 2.9486 |

